# Coevolution between life-history and metabolic rate depends on ontogenetic stage

**DOI:** 10.1101/705707

**Authors:** Will Sowersby, Sergey Morozov, Simon Eckerström-Liedholm, Philipp Lehmann, Piotr K. Rowiński, Joacim Näslund, Alejandro Gonzalez-Voyer, Björn Rogell

## Abstract

Metabolic rate is considered to determine the energetic investment placed into life-history traits, regulating the speed of an organism’s life-cycle and forming the so called “pace-of-life”. However, how metabolic rate and life-history traits co-evolve remains unclear. For instance, the energetic demands of life-history traits, including the number of energy allocation trade-offs, is unlikely to remain constant over ontogeny. Therefore, the predicted coevolution between metabolic rate and life-history could be specific to particular ontogenetic stages, rather than a stable property of an organism. Here, we test the ontogenetic dependency of the coevolution between metabolic rate and the pace of life-history, under strictly standardized conditions using 30 species of killifish, which are either annual species adapted to ephemeral pools or non-annual species inhabiting more permanent waterbodies. Standard metabolic rates were estimated at three ontogenetic stages, together with relevant life-history traits, i.e. growth (juveniles), maturity (young adults), and reproductive rate (reproductive adults). Life-history traits largely followed predicted pace-of-life patterns, with overall faster/higher rates in annual species. The divergences in life-history traits across species tended to increase over ontogeny, being smallest during juvenile growth and largest in reproductive adults. However, associations between life-history strategy and metabolic rate followed a reversed pattern, being strongest in juveniles, but lowest in reproductive adults. Our results are concordant with the number of energetic trade-offs increasing over ontogeny, which results in a stronger covariation between physiology and life-history traits earlier in ontogeny.

## Introduction

An organism’s position along the fast-slow life-history continuum is typically linked to differential investment into energetically costly traits (e.g. growth, development, reproductive rate; 1–5). Metabolic rate is considered to be an important functional trait in determining the rate at which resources are converted into the energy available for life-history traits (6,7), and the maintenance of somatic tissues (8). While presumably not under direct selection (9), evolutionary shifts in metabolic rate are thought to follow concomitant changes in the pace of life-histories (10). However, despite clear theoretical expectations, empirical evidence for the co-evolution between metabolic rate and the pace of life-history remains ambiguous, with mixed results from both experimental and comparative studies (e.g. 10–12).

Life-history traits and metabolic rate have commonly been measured on individuals sampled from the wild, rather than individuals reared under standardised experimental settings (10). This lack of standardisation has potentially obscured the coevolutionary associations between the pace of life-history and metabolism, due to plasticity in either life-history traits or metabolic rate (life-history traits: 13; metabolic rates: 14,15). Furthermore, the relationship between metabolic rate and life-history is likely to vary over the course of an organism’s life-cycle, rather than remaining a stable property of an organism (13). For instance, the more energetically costly elements of a life-history, which vary substantially across species (15,16), should be most prominent at particular ontogenetic stages (e.g. juvenile - growth; adult - reproduction). In contrast, for a stable relationship between life-history and metabolism to occur throughout ontogeny, the amount of energy invested into somatic growth should be equal to that later invested into reproduction, resulting in a constant energy allocation toward these relatively costly life-history traits. However, a fast rate of growth does not necessarily lead to a high reproductive rate, and vice versa (i.e. the pace of a life-history will not predict all variation in a system; 1). Partly, this is due to an organism not being a tightly integrated entity, but rather consisting of quasi-independent parts, which are tightly integrated (16). This modularity reduces the probability for trade-offs to influence evolutionary change (17), as a finite amount of energy being allocated among processes (e.g. homeostasis), results in the number of trade-offs (18) typically increasing over ontogeny (19). Therefore, this increasing number of trade-offs may mask associations between life-history traits and metabolic rate, and result in the latter being influenced by a complex co-regulation between life-history characters and physiology that can change over an organism’s lifespan. As a consequence, we argue that there is a need to integrate ontogeny into the pace-of-life framework, as the coevolution between metabolic rate and the pace of life-history could remain undetected, if changes in energy allocation towards different life-history traits over ontogeny are not properly considered.

Here, we test the association between metabolic rate and life-history traits (somatic growth rate, maturity and reproductive rate) across different ontogenetic stages (juveniles, young non-reproductive adults, and reproducing adults). We do this by conducting a large-scale comparative, common garden study of 30 killifish species from the Aplocheiloidei suborder, focusing on rates of growth, reproduction, maturity, and standard metabolic rate (SMR). Within this clade there has been several independent evolutionary transitions between annual and non-annual life-history strategies, where annual species have evolved egg-stages capable of entering embryonic diapause to adapt to the regular desiccation of ephemeral aquatic habitats (20). In contrast, non-annual killifishes inhabit more permanent, stable habitats, allowing these species to live and breed over comparatively longer time-scales. Adaptations to time-constrained environments, such as ephemeral habitats, typically include correlated selection on increased rates of growth (21), quicker development periods (21,22); including in killifishes (23), as well as higher reproductive rates (24). Moreover, annual killifishes have very short life-spans in comparison to other similar vertebrates, which is likely a consequence of reduced selection on later-life fitness (25,26,23,27–30). As such, we expect the independent evolutionary transitions between annual and non-annual life-history strategies in killifishes (20) to be accompanied by the evolution of life-history traits along the fast-slow life-history continuum. Hence, killifishes present an unparalleled system for conducting comparative analyses on organisms which are ecologically similar, yet potentially diverge substantially in regard to the pace of life-history traits.

In killifishes, we purposely investigate life-history traits that are directly linked to biosynthesis, and are therefore governed by an allocated energetic budget, predicted to be positively correlated with metabolic rate (31–36). We use a strictly standardized common garden experimental set-up, as plasticity in both metabolic rate and life-history traits has been suggested as a potential source of ambiguity in the empirical evidence previously collected on the pace of life-histories (10). We predict that life-history traits and metabolic rate in killifishes should evolve in a correlated manner, and that annual species should on average exhibit faster life-history traits and have a higher metabolic rate, compared to non-annual species. Furthermore, if metabolic rate evolves to fuel life-histories, we predict that the maximum divergence between comparable species with fast and slow life-histories will coincide with the ontogenetic time point when corresponding life-history trait divergences are also at their maximum.

## Methods

### Study system

Life-history strategy (annual or non-annual) is characterised in killifishes according to the presence or absence of eggs capable of entering embryonic diapausing (20). Here, we reared 30 species of killifish, 13 non-annual and 17 annual species (see Supplementary Table 1 for the full species list), which we selected based on their phylogenetic position (20), and to represent multiple independent evolutionary transitions to an annual life-history. Diapausing eggs have both lower expressions of growth hormones (37,38) and lower metabolic rates (39,40) compared to directly developing eggs, implying that the diapause stage is an adaptation to ephemeral habitats that regularly desiccate for extended periods, and is not mechanistically linked to traits related to the pace-of-life.

Fish were housed under laboratory conditions (average 24.3°C; 12-hour day:night cycle), and were fed newly hatched *Artemia*, supplemented with frozen bloodworms when they reached adulthood, to satiation three times daily (once daily during weekends). All individuals were hatched from eggs under our laboratory conditions, with eggs either produced from our own stock populations, or sourced from dedicated aquarists. Fry were initially housed in small plastic containers (9 × 9 × 9 cm) and were moved to bigger aquaria as they grew larger (13 L; furnished with gravel, clay pots and bundles of wool yarn). All fish were initially raised in solitary conditions, but after sexual maturity a subset were housed as pairs, or trios (1 male and 2 females, to reduce male aggression) and allowed to breed, with the remainder kept in isolation and unable to breed, until the end of the experimental period.

### Growth rate

We measured the growth rate of 29 species (*N*_annual_ = 16; *N*_non-annual_ = 13; *N* = 400 individuals; Supplementary Table 1), during the linear juvenile growth phase, by photographing each individual every 7 to 10 days, from hatching until an age of ca. 3 months or sexual maturity. Standard body length was measured from these photographs using the software ImageJ (41). As the level of replication in our analysis was at the individual, any imprecisions in measurements due to minor distortions in images could have substantial inferential impact. Therefore, to avoid incorporating high amounts of noise in the data, we fitted Gompertz growth models to each individual, and removed outliers that had absolute values of residuals >0.11. Outlier removal was script-based and hence blind to species identity. Using data on length, with outliers removed, we fitted a linear model for each individual, in the interval between 10 and 60 days of age, and used the slope of this regression as a measure of the growth rate of the individual, which was predominantly linear (see supplementary material). We also assessed growth as the Gompertz-parameter μ; however, as growth was linear (e.g. most species had not reached the plateau of the growth phase) during the measurement period, μ was estimated outside the range of the data in more than half of the individuals. Hence, here we use the slope of a linear regression of time on length as a proxy for individual growth, while analysis of µ yielded congruent results (results not presented).

### Maturity

Killifishes are sexually dimorphic in both shape and colour (27; Sowersby et al. under review), which we exploited to assess developmental rates. Specifically, as juveniles are typically more similar in appearance to females than males, we noted the time point (in days since hatching) at which we were able to determine if an individual was a male (based on the appearance of species-specific male colour patterns) using the photos taken weekly for growth estimation. Our measure of developmental rate was hence sex-specific and assumed that both females and males mature at similar time points. This should be a valid assumption, given that the life-history evolution of these fishes is likely to be predominantly determined by time-constrained environmental conditions, and therefore similar for both sexes. Furthermore, in other studies we have dissected killifishes at various time points in their life-cycle and have found that sex identification based on visual inspection always aligns with the correct male or female reproductive tissue being observed during dissection (Sowersby et al. under review). Our predictions were that annual species would have higher values (i.e. higher rates) in all measured traits, compared to non-annual species. Therefore, for ease of interpretation, we subtracted each individual’s time to maturity observation from the overall mean time to maturity, across all species, creating an index where small values correspond to fast development and large values with slow development time (i.e. an additive inversion of the data, henceforth “rate of maturity”). The analysis was performed on 113 individuals from 22 species (*N*_annual_ = 12; *N*_non-annual_ = 10), where the sample size ranged from 1 to 10 individuals per species (median: 4.5; Supplementary Table 1). As killifishes sometimes show significant bias in sex-ratios (Sowersby et al. under review), sample sizes differed among species, depending on the number of males available.

### Reproduction

We used previously estimated reproductive rates (24). Briefly, for 19 species (*N*_annual_ = 11; *N*_non-annual_ = 8; Supplementary Table 1), of which 16 overlap with species used in the growth measurements, we estimated reproductive rates by counting the number of eggs deposited by each female per month. Females that did not reproduce during this month, where considered to be reproductively inactive, and were excluded from the data.

### Standard Metabolic Rate

Oxygen consumption was measured over time using an intermittent-flow respirometry setup (Loligo Systems, Viborg, Denmark), set at 24°C, under a 12:12 hour day:night regime (light: 07:00 – 19:00). Specifically, we measured oxygen uptake rates (as a proxy for estimated SMR) at three different biologically relevant ontological time points (juveniles: *N* = 187, from 13 slow-living and 16 fast-living species; young adults *N* = 141, from 13 slow-living and 17 fast-living species; reproducing adults: *N* = 223, from 10 slow-living and 10 fast-living species; see Supplementary Table 2 for details). Prior to the measurements, fish were fasted for 15 hours (8 hours in home tanks, 7 hours during acclimation in the respirometry chambers) and weighed (prior to acclimation). Trials were run for ca. 17 h, overnight, starting approximately at 17:00. Oxygen consumption was measured in total darkness, in 30-minute cycles (between 00:00 - 05:00, when fish were likely to be least active), by estimating the slope of the decreasing oxygen concentration over time. Out of ten slopes obtained per individual, the three lowest-valued slopes (the majority being R^2^ > 0.9) were retained for further analysis. Of these, all analyses were performed on the lowest of these values. The lowest estimate of oxygen consumption was strongly correlated to the mean of the 3 lowest values (r = 0.998), and analysis on the mean of the three lowest values yielded highly congruent results. The decrease in oxygen consumption over time was highly linear (typically R^2^ > 0.98), suggesting that fish were consistently inactive during this period. The rate of background respiration (due to the presence of microorganisms in the respirometry set-up) was accounted for by taking blank tests (i.e. with no fish) before and after SMR measurements, and subtracting the extrapolated value from the total oxygen consumption. The respirometry analysis, including correction for background respiration, was conducted using the R package FishResp (42). We performed a Cook’s D outlier analysis on a model with log_10_ oxygen consumption as response, and log_10_ mass as a covariate, including species as a random effect (fit using restricted maximum likelihood in the R-library lme4), and the most deviating observations (i.e. the 5% quantile; 17 observations) were removed. In a subset of analyses we used residual metabolic rate, which was calculated as the residuals from a regression on log_10_ oxygen consumption as the response, and log_10_ mass as a covariate, log_10_ oxygen consumption as a response, log_10_ mass, ontogenetic stage, and their interaction as explanatory variables.

### Phylogeny

In order to control for phylogenetic non-independence of data points, we included information on shared ancestry based on a dated phylogeny (20). We added missing taxa to the dated phylogeny by utilizing other previously published phylogenies (see supporting information), using the add.species.to.genus and bind.tip functions in the R package phytools (43). The resulting final phylogeny was included as a random factor in all Bayesian linear mixed models (see below).

### Statistical analysis

We aimed to test how life-history traits and metabolic rate co-evolved across species, as well as whether the association between life-history and metabolic rate is dependent on ontogenetic stages. Given the difficulty in disentangling correlations between species means into a component caused by phylogenetic signal and a component caused by evolutionary processes independent from ancestry, we analysed our data utilizing our a priori sampling regime, i.e. species that were sampled from repeated, independent evolutionary transitions between two states: absence-presence of diapausing eggs (a specific adaptation to ephemeral habitats). By employing phylogenetically informed sampling we were able to obtain a number of phylogenetically independent contrasts, between species that differed in the pace of life-history traits associated with adaptations to ephemeral or permanent habitats. However, as outlined in the introduction, theory predicts correlations among species, inferences from our models therefore assume that any divergences across annual and non-annual life-history groups arise due to correlations among species. In order to validate this assumption, we also analysed species-level correlations. Further, as theory predicts that metabolic rate evolves as a correlated response to the energy requirements of particular life-history traits, we should expect that standardised differences in life-histories will be comparable to standardised differences in metabolic rates. To assess this possibility, we compared standardised contrasts in life-history traits, across annual and non-annual species, with standardized contrasts in the corresponding metabolic rates, at the matching ontogenetic stage.

### Contrasts between annual and non-annual life-history strategy

We first tested the effect of life-history on rates of growth, maturity and reproduction, as response variables in a multivariate model, with the trait specific means, life-history strategy (annual or non-annual) and their interaction as fixed effects. Species and phylogeny were added as random effects, as well as separate residual variances for each response variable.

Then, to test if metabolic rate differed between annual and non-annual species, we fit the lowest measured value of oxygen consumption (i.e. standard metabolic rate) per individual as a log_10_ transformed response variable, with log_10_ transformed body size, the presence or absence of diapausing eggs (i.e. annual or non-annual), ontogenetic stage, and the interaction between ontogenetic stage and presence or absence of diapausing eggs added as fixed effects, species, phylogeny, and the interaction variance of species and ontogenetic stage were added as random effects. We did not explicitly focus on any sex differences, as the life-history trait variables we measured were either independent of sex (e.g. juvenile growth rate), or were purposely defined by only one sex (e.g. we used the secondary sexual traits of males to assess sexual maturity and the number of eggs laid by females were used as a proxy for reproductive rates, see (24). However, it has been previously suggested that any coevolution between metabolic rate and life-history could be sex-specific (44,45). Therefore, we tested this hypothesis by modelling log_10_ transformed metabolic rate as a response variable, and log_10_ transformed body size, the presence/absence of diapausing eggs, ontogenetic stage, and sex as fixed effects. No interactions among the fixed effects were significant, and were hence not included. Species, phylogeny, and the interaction variance of species and ontogenetic stage were added as random effects.

### Correlations among traits and ontogenetic stages

To assess species level correlations between metabolic rate, life-history traits, and ontogenetic stages, we analysed *i)* a multivariate model with rates of growth, development and reproduction as responses), *ii*) specific life-history traits (in total three bivariate models, growth - juvenile; maturity rate - young adults; reproductive rate - reproductive adults), *iii*) a univariate model of residual metabolic rate as a response variable and ontogeny included as a fixed effect. In these models, we fitted the full covariance matrices associated with species-specific and phylogenetic effects. We examined the amount of variation explained by species using intra class correlations. The putative species-level correlations among traits, were calculated from the species-level covariance matrix, calculated as the sum of the phylogenetic covariance matrix, and the species-specific covariance matrix (46).

### Standardized contrasts across annual and non-annual species

To compare contrasts across life-history strategies (i.e. annual and non-annual) based on traits with different means and variances, we calculated effect sizes (Hedge’s D) from the posterior distributions. For life-history traits, the means were extracted from the multivariate model testing the effect of annual and non-annual strategy on measured life-history traits. For metabolic rate, means and variances were extracted from a model where residual metabolic rate was fit as the response variable, and ontogenetic stage, life-history and their interaction were fit as fixed effects, with species and phylogeny fit as random effects.

All models were fit within a (multivariate) Bayesian phylogenetic mixed model framework, using the R package MCMCglmm (47), with flat priors for the fixed effects, and correlations. Three chains were fit to each model, and the posterior modes were fused across these models. Within chains, autocorrelations were in the interval >-0.1 and <0.1, and the Gelman diagnostic across the three replicate chains was always <1.2, both suggesting that the Bayesian models converged. Flat priors were used for the fixed effects, and inverse Wishart priors, flat for the correlation, were used for the random effects.

## Results

### Life-history traits

In congruence with our predictions, annual species exhibited an overall faster pace-of-life, with faster rates of growth, maturity rates, and higher reproductive rates (see 24), in comparison to non-annual species (Table 1; Figure 1). All life-history traits exhibited variation explained significantly by species, where species explained 77.9% of the variation in growth rates (95% CI: 67.4-86.6), 64.6% in maturity rate (i.e. inversed time to maturity; 95% CI: 47.4-78.8), and 82.2% in reproductive rate (95% CI: 61.8-92.7) (Supplementary Table 2). Life-history traits further exhibited significant correlations across species (Figure 2b). Importantly, sample sizes for growth rates were higher than both maturity and reproductive rates, meaning that growth data may exhibit higher precision for species-specific estimates. However, growth rates had the lowest intra-class correlation for species, which indicates that the other two life-history rates had larger underlying biological effect sizes. Hence, it is unlikely that any results were driven by varying statistical power across the three traits.

**Table 1:** Results of the Bayesian phylogenetic mixed model testing how the pace of life-history traits (growth, maturity and reproductive rates) depends on life-history strategy (annual or non-annual), with life-history strategy and the interactions between life-history traits and life-history strategy as fixed effects. The model was run with the specific species and phylogeny used in each life-history trait as random effects. Where, “species” signifies the variance explained by species, “phylogeny” signifies the variance explained by the phylogeny, and “residual variance” is the variance that is not explained by the mode. Lower and upper CIs represent 95 % credible intervals. The table is structured according to the standard output from R.

| Parameter | Estimate | Lower CI | Upper CI | P<sub>MCMC</sub> |
| --- | --- | --- | --- | --- |
| *Fixed Effects* |  |  |  |  |
| Growth Rate | 0.0355 | 0.0268 | 0.0453 | 0.000333 |
| Maturation Rate | 6.78 | -4.51 | 18.9 | 0.177 |
| Reproductive Rate | 3.93 | 2.94 | 4.78 | 0.000333 |
| Life-History | -0.0133 | -0.0213 | -0.00284 | 0.0153 |
| Maturation Rate : Life-History (Non-Annual) | -23.6 | -34.2 | -12.9 | 0.000333 |
| Reproductive Rate : Life-History (Non-Annual) | -1.93 | -2.92 | -0.769 | 0.000667 |
| *Random Effects* |  |  |  |  |
| Growth Rate (Phylogeny) | $7.93 \times 10^{-07}$ | $3.17 \times 10^{-11}$ | 0.000254 | - |
| Maturation Rate (Phylogeny) | 2.28 | $6.59 \times 10^{-07}$ | 317 | - |
| Reproductive Rate (Phylogeny) | 0.0157 | $2.05 \times 10^{-07}$ | 1.56 | - |
| Growth Rate (Species) | $8.41 \times 10^{-07}$ | $4.60 \times 10^{-11}$ | 0.000154 | - |
| Maturation Rate (Species) | 0.863 | $9.08 \times 10^{-06}$ | 174 | - |
| Reproductive Rate (Species) | 0.0048 | $1.30 \times 10^{-07}$ | 0.892 | - |
| Growth (Residual Variance) | 0.000122 | $9.87 \times 10^{-05}$ | 0.00014 | - |
| Maturation Rate (Residual Variance) | 139 | 108 | 199 | - |
| Reproductive Rate (Residual Variance) | 0.274 | 0.163 | 0.502 | - |

**Figure 1.**
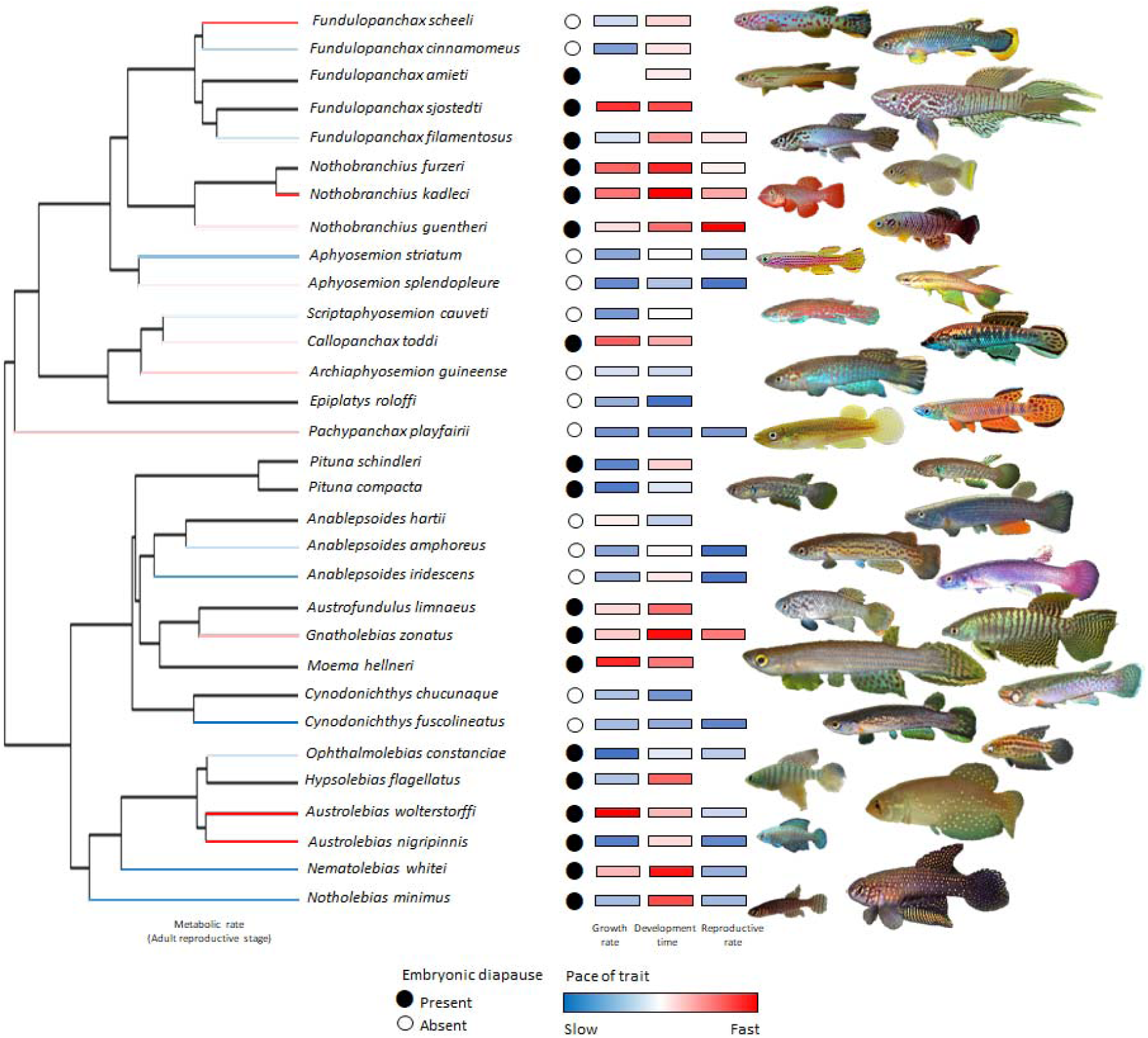
Updated phylogenetic tree (utilizing Furness et al. 2015) of the species used in the study. Adult breeding stage residual standard metabolic rate (SMR was highly correlated across ontogenetic stages) is displayed on the tips of the tree. The intensity of red (fast) and blue (slow) colour represents the mean pace of measured life-history traits (growth rate, development time, and reproductive rate), per species.

**Figure 2.**
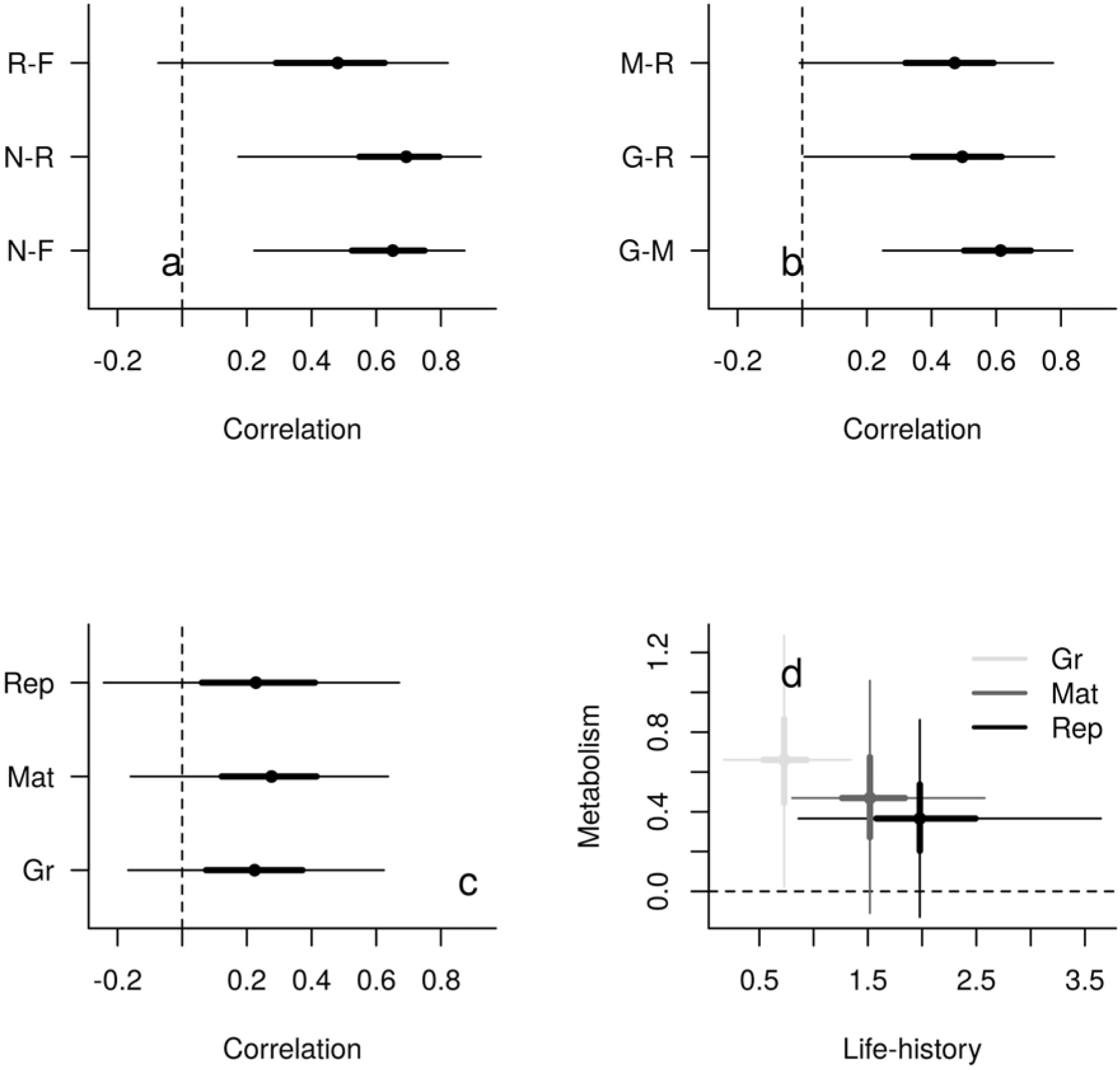
Correlations between a) species level metabolic rates across ontogenetic stages, where R = reproductive adults, N = non-reproductive adults, and F = juveniles. b) The estimated species level correlations between different life-history traits where G = growth, M = time to maturity, and R = reproduction. c) The estimated species level correlations between metabolic rate and reproduction (Rep), maturity rate (Mat), and growth (Gr). d) Effect sizes (Hedges d) of divergences following adaptations to ephemeral environments for life-history traits and ontogeny specific metabolic rates, for juvenile-growth, non-reproductive adults-maturity rate and reproductive adults-reproductive rate. In all plots, the dot is the median of the posterior distribution, narrow lines denote the 95% credibility intervals, and thick lines indicate the 50% credibility intervals.

### Metabolic rates

We found that for a given body size, metabolic rate differed between annual and non-annual species, with annual species having overall higher metabolic rates than non-annual species [beta = -0.081 (95% CI: -0.16 --0.018), P = 0.02, Table 2]. Further, we found that metabolic rates, in general, decreased over ontogeny, being highest in fry and lowest in reproductive adults (Table 2). The allometric slope for metabolic rate was 0.88 (95% CI 0.83 - 0.92) and we found no significant differences in the allometric slope of metabolic rate between the two life-history strategies or between the ontogenetic stages. We found no significant differences among the sexes in metabolic rate (Supplementary Table 3).

**Table 2:** Results of the Bayesian phylogenetic mixed model on standard metabolic rate, with ontogenetic stage (juvenile, non-reproducing adult, reproducing adult), life-history strategy (annual or non-annual), fish mass (log10 transformed) and the interaction between ontogenetic stage and life-history as fixed effects. The model was run with species, phylogeny and the interaction variance of ontogenetic stage and species as random effects. Where, “species” signifies the variance explained by species, “phylogeny” signifies the variance explained by the phylogeny, “ontogenetic stage : species (interaction variance)” represents the variance explained by ontogenetic stage and species and “Residual Variance” is the variance that is not explained by the model. Lower and upper CIs represent 95 % credible intervals.

| Parameter | Estimate | Lower CI | Upper CI | P<sub>MCMC</sub> |
| --- | --- | --- | --- | --- |
| *Fixed Effects* |  |  |  |  |
| (Intercept) | -0.812 | -0.895 | -0.726 | 0.000333 |
| Ontogenetic Stage (Non-Reproducing Adult) | -0.109 | -0.171 | -0.0598 | 0.000667 |
| Ontogenetic Stage (Reproducing Adult) | -0.15 | -0.218 | -0.103 | 0.000333 |
| Life-History (Non-Annual) | -0.105 | -0.176 | -0.0198 | 0.0147 |
| Fish Mass | 0.884 | 0.828 | 0.922 | 0.000333 |
| Ontogenetic Stage (Non-Reproducing Adult) : Life-History (Non-Annual) | 0.0242 | -0.0362 | 0.0994 | 0.415 |
| Ontogenetic Stage (Reproducing Adult) : Life-History (Non-Annual) | 0.0202 | -0.0419 | 0.11 | 0.336 |
| *Random Effects* |  |  |  |  |
| Species | $4.69 \times 10^{-05}$ | $7.01 \times 10^{-10}$ | 0.00799 | - |
| Phylogeny | $4.51 \times 10^{-05}$ | $8.41 \times 10^{-09}$ | 0.0134 | - |
| Ontogenetic Stage : Species | 0.00206 | 0.000665 | 0.00437 | - |
| Residual Variance | 0.0111 | 0.00962 | 0.0123 | - |

When assessing species effects in the different ontogenetic stages, we found significant species effects across all stages of ontogeny, but the variation explained by species decreased over ontogeny. Species explained 63.2 % (95% CI: 44.9 – 78.6) of the variation in juveniles, 53.8% (95% CI: 37.5 – 69.8) in non-reproducing young adults, and 31.3% (95% CI: 14.8 – 53.9) in reproductive adults, due to lower among-species variation at later stages of ontogeny (Supplementary Table 4). Further, species-specific metabolic rates were strongly correlated across ontogenetic stages, suggesting that the clustering into high vs low metabolic rates is a species-specific property that is stable over ontogeny (Figure 2a). Species level correlation in metabolic rate was 64.5 between juveniles and young adults (95% CI: 20.5 – 86.8), 47.7 between juveniles and reproductive adults (95% CI: -9.25 – 81.9), and 68.4 between young adults and reproductive adults (95% CI: 16.6 – 92) (Supplementary Table 4).

### Connections between life-history and metabolic rate

When examining the contrasts between annual and non-annual species in metabolic rate specific to the three life-history stages (Supplementary Table 4), to contrasts in life-history traits expressed at these ontogenetic stages, we found trends towards diminishing contrasts in metabolic rate over ontogeny, but increased contrasts in life-histories, where growth rate had the smallest difference between annual and non-annual species and reproductive rate had the largest difference (Figure 3d). Correlations between life-history traits and metabolic rate were in general weak, and while none were significantly different from 0, growth rate had a stronger correlation than development time and reproduction (Figure 3a; Supplementary Tables 5-7).

**Figure 3.**
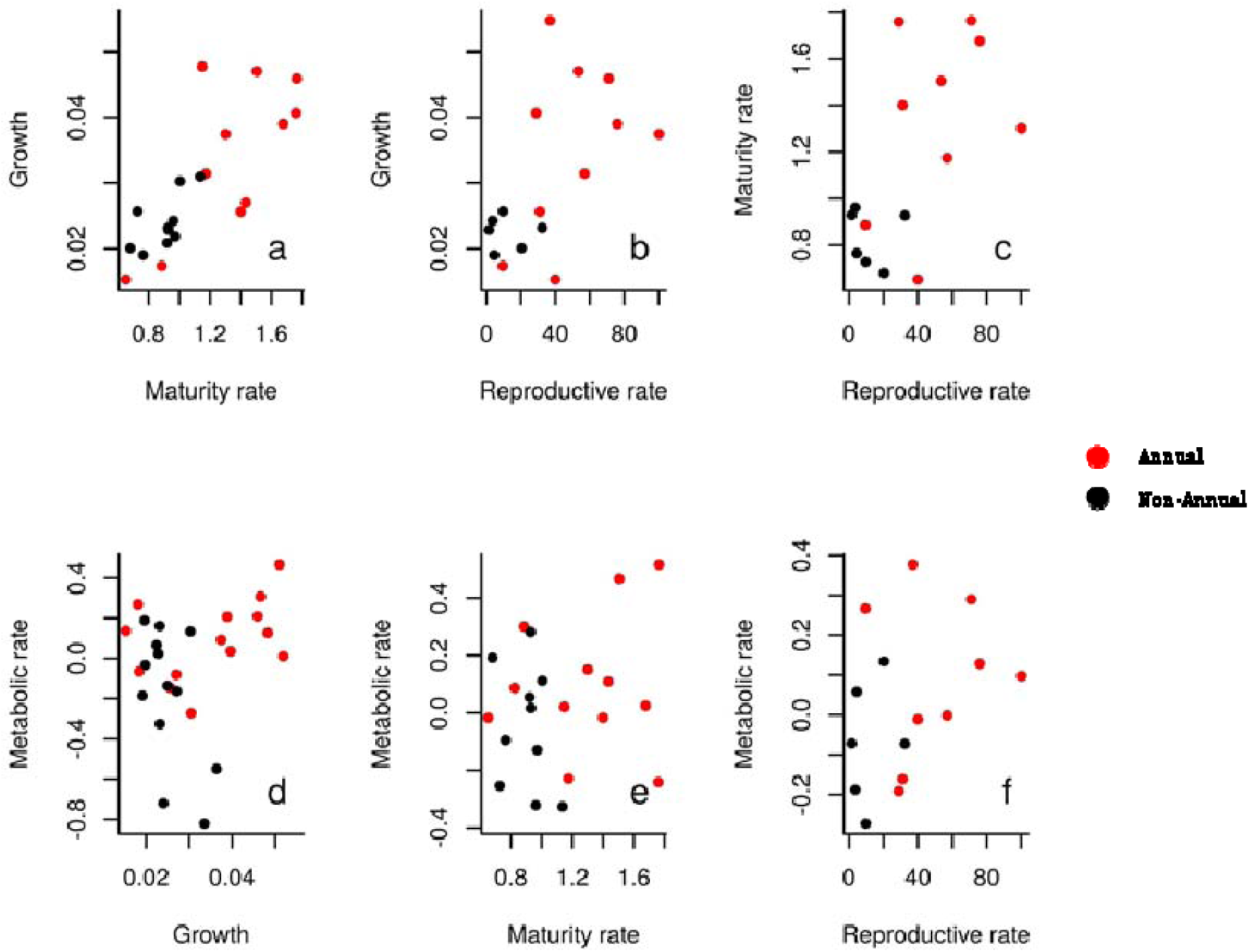
Scatterplots between a) rates of growth and maturity. b) rates of growth and reproduction. c) rates of maturity and reproduction. d) growth rate and metabolic rate for juveniles, e) maturity rate and metabolic rate for non-reproductive adults, and f) reproductive rates and metabolic rates of reproductive adults. In all plots, red dots indicate annual species, and black dots non-annual species.

## Discussion

In accordance with the predictions made by pace-of-life theory, we found that rates of life-history traits were correlated across species. Specifically, we found that annual species had significantly faster rates of growth, maturity, and reproduction, in comparison to non-annual species. The divergences between life-history strategies (annual and non-annual) tended to increase over ontogeny, being smallest during juvenile growth, and largest in reproductive adults. In general, metabolic rate was higher in annual fishes, and followed a similar pattern with species means correlating positively over the three ontogenetic stages. Interestingly, we found that the rank order of metabolic rate across species was relatively stable over ontogeny, implying that placement along the axis of low to high metabolic rate can be considered a species-specific trait.

However, while we found no significant interactions between life-history and ontogenetic stage, we found that associations between metabolic rate and life-history strategy were strongest at earlier ontogenetic stages (i.e. at the juvenile stage), and tended to decrease over ontogeny. Rather than being a straightforward energetic trade-off between investment into growth or reproduction, our results suggest a more complex relationship between metabolic rate and life-history, where energetic allocations likely change over the course of an organism’s life-cycle. Furthermore, as our analyses were phylogenetically controlled, our results suggest that pace-of-life is sustained across genetically diverged species, including independent evolutionary transitions, e.g. suggesting parallel evolution.

### Life-histories co-evolve as predicted by pace-of-life theory

To maximise reproductive success, life-history traits are predicted to evolve in response to different biotic and abiotic environmental factors (15). According to life-history theory, species with high rates of extrinsic mortality are typically selected to emphasise current over future reproductive events, which often results in the co-evolution of life-history traits in the same direction, along a fast-slow continuum (15). We found that key life-history traits did indeed correlate in the direction predicted by pace-of-life theory. Specifically, annual killifish species, which are adapted to time-constrained ephemeral habitats, exhibited significantly faster rates of all measured life-history traits, compared to non-annual species. While evolution under differential mortality rates has been identified as a key driver of the pace of life-histories (15,48), annual and non-annual killifishes do often co-occur, for example in flood plain areas (20; Sowersby et al. under review). Hence, we acknowledge that adult annual killifish could possibly escape the desiccation of ephemeral habitats by seasonally migrating to more permanent water-bodies, like some non-annual species (49). However, the strong divergences we observed in the pace of life-histories among the two groups suggests that these migrations may not occur, or do not occur at evolutionarily significant frequencies. In addition to having eggs capable of entering diapause, annual killifishes are also known to age rapidly compared to other similar vertebrate species, which is highly unlikely to be an adaptive trait if these fishes are migrating away from ephemeral environments (26). Therefore, adaptations to ephemeral, time-constrained environments, appear to have played a key role in shaping the evolution of the pace of life-history, in this clade of fishes.

### A faster life-history is associated with a higher metabolic rate

One hypothesis that has received substantial interest stipulates that life-history strategies characterised by high rates of growth and reproduction, should have co-evolved corresponding physiological mechanisms to fuel these energetically demanding processes (4). In this context, the pace-of-life hypothesis has been proposed as a framework explaining the expected coevolution of metabolic rate and life-history strategies (3,50,6,10). However, the empirical evidence for this relationship between metabolic rate and the pace of life-histories has remained weak (13; including in killifishes, Eckerström-Liedholm et al. under review). One plausible explanation for this disparity between theoretical predictions and empirical results may be because both metabolic rate and life-history traits have considerable plasticity (51,14,52,53), meaning any coevolutionary associations are potentially distorted by environmental effects (10,54; see 13). Here, we controlled for potential confounds generated by environmental effects, by employing a common garden approach with strictly standardised environmental conditions. Under these conditions we found that metabolic rates were indeed higher in species with an annual life-history, compared to non-annual species. Further, metabolic rate generally decreased over ontogeny, which has been attributed to a decrease in the relative size of metabolically costly tissues (e.g. the liver) over ontogeny (55). For example, a decreasing proportion of organs with high mass-specific metabolic rates throughout ontogeny provides a proximate mechanistic explanation for ontogenetic declines in metabolic rate (56–58). Although metabolic rate often displays plasticity and decreases over ontogeny, we found that species level correlations between metabolic rate, measured at three different ontogenetic stages, were positive and rather strong. This implies that species level rank orders of metabolic rate remained rather consistent, which is in congruence with previous research on metabolic allometries across fishes (59).

The total energy budget of an organism is distributed amongst key functions, such as activity, biosynthesis (growth and reproduction), and somatic maintenance (59). As resources are typically finite, energy used for one function diminishes that available for another, creating energetic conflicts. For instance, both theoretical and empirical evidence indicates that organisms with fast rates of growth and reproduction have shorter lifespans, suggesting that fast and slow-living organisms invest different amounts of their energy budget into the maintenance of somatic tissues (59). Indeed, annual killifishes, which we found to have generally fast rates of growth and reproduction, do have short lifespans, with some *Nothobranchius* species having among the shortest lifespans recorded for any vertebrate (60). Therefore, it is possible that slow-living non-annual killifish species could have overall similar metabolic requirements as annual fast-living killifishes, if non-annual species invest a greater amount of energy into somatic maintenance. However, we found that annual species, which had faster rates of both growth and reproduction, also had significantly higher metabolic rates. This pattern indicates an association between the life-history traits directly involved in biosynthesis (e.g. growth and reproduction) and energetic demands, which is presumably in excess of the energy invested by non-annual species into somatic maintenance. Our results are hence largely congruent with Pettersen et al. (61), who found that bryozoans with a higher metabolic rate have shorter developmental times and life-spans, in contrast to bryozoans with a lower metabolic rate.

### Across species, pace-of-life patterns are most apparent early in ontogeny, when difference in metabolic rate are the smallest

When assessing standardised differences in life-history traits, we found that divergences across annual and non-annual species increased over ontogeny. This is not an unexpected pattern, as evolutionary trajectories frequently occur through changes in the timing and the rate of developmental events, leading to an accumulation of divergence throughout life (62,63). As a consequence, evolutionary divergence has been found to increase over ontogenetic development (64,65). However, surprisingly, patterns of pace-of-life have not typically (if at all) been assessed over ontogeny, and ours is the first to do so at a macro-evolutionary scale.

If differences in metabolic rate evolve as a correlated response to selection on key life-history traits (9), we would expect that differences in metabolic rate between life-history strategies would become increasingly evident during the later stages of ontogeny. However, we found that associations between life-history strategy and metabolic rate followed a reversed pattern, being significantly different in juveniles, but tending to decrease over ontogeny (i.e. being lowest in reproductive adults). More specifically, standardized divergences (between annual and non-annual species) in metabolic rate and growth, were of roughly the same magnitude in juveniles, while divergence in reproductive rate was approximately five times higher than the divergence in metabolic rate in adults. This pattern suggests that the number of energetic trade-offs within an organism increases over ontogeny (19), with the energetic trade-offs that exist earlier in development potentially involving lower modular complexity. If the number of potential energetic trade-offs does increase over time within an organism, pace-of-life patterns may be most apparent earlier in ontogeny. Overall, our results suggest that the relationship between metabolic rate and life-history is likely to be influenced by a complex interaction between life-history characters and physiology, which is modulated over ontogeny.

## Conclusion

In conclusion, we found that annual killifishes that are adapted to ephemeral environments had overall faster life-histories, both in terms of life-history traits and in associated changes in metabolic rate. This is in comparison to non-annual killifishes that are adapted to more stable, constant environments. We had predicted that differences between fast and slow life-history strategies, in regard to the association between life-history traits and physiology, would be most apparent at distinct points during ontogeny. Indeed, we found that associations between life-history and metabolic rate were higher during periods of juvenile growth. Our results show that the covariance between metabolism and life-history traits can change over ontogeny, likely due to an increase in the number of trade-off components as an organism develops into a reproductive stage.

