## Supporting Information for "Coevolution between life-history and metabolic rate depends on ontogenetic stage"

**Supplementary Table 1**

| Species | Life-History | Growth Rate | Maturity | Reproductive Rate | Juvenile SMR | Adult SMR (Non-Reproducing) | Adult SMR (Reproducing) |
| --- | --- | --- | --- | --- | --- | --- | --- |
| *Anablepsoides hartii* | Non-Annual | 8 | 2 | NA | 3 | 2 | NA |
| *Aphyosemion splendopleure* | Non-Annual | 18 | 6 | 4 | 8 | 8 | 7 |
| *Aphyosemion striatum* | Non-Annual | 10 | 9 | 9 | 9 | 9 | 16 |
| *Archiaphyosemion guineense* | Non-Annual | 3 | 2 | NA | 3 | 3 | 4 |
| *Austrofundulus limnaeus* | Annual | 8 | 1 | NA | 4 | 5 | NA |
| *Austrolebias nigripinnis* | Annual | 14 | 4 | 3 | 3 | 2 | 1 |
| *Austrolebias wolterstorffi* | Annual | 12 | 2 | 3 | NA | 3 | 6 |
| *Callopanchax toddi* | Annual | 14 | 4 | NA | 10 | 9 | 14 |
| *Cynodonichthys chucunaque* | Non-Annual | 17 | 2 | NA | 4 | 3 | NA |
| *Epiplatys roloffi* | Non-Annual | 12 | 2 | NA | 6 | 4 | NA |
| *Fundulopanchax amieti* | Annual | NA | 2 | NA | 5 | 4 | NA |
| *Fundulopanchax cinnamomeus* | Non-Annual | 9 | 5 | NA | 6 | 6 | 15 |
| *Fundulopanchax filamentosus* | Annual | 27 | 5 | 1 | 8 | 6 | 17 |
| *Fundulopanchax scheeli* | Non-Annual | 14 | 8 | NA | 9 | 7 | 17 |
| *Fundulopanchax sjoestedti* | Annual | 8 | 2 | NA | 5 | 5 | NA |
| *Gnatolebias zonatus* | Annual | 25 | 4 | 1 | 9 | 3 | 17 |
| *Hypsolebias flagellatus* | Annual | 12 | 2 | NA | 9 | 4 | NA |
| *Moema hellneri* | Annual | 12 | 2 | NA | 4 | 1 | NA |
| *Nematolebias whitei* | Annual | 15 | 10 | 9 | 8 | 5 | 15 |
| *Nothobranchius furzeri* | Annual | 6 | 2 | 2 | NA | 4 | NA |
| *Nothobranchius guentheri* | Annual | 14 | 5 | 4 | 10 | 9 | 17 |
| *Nothobranchius kadleci* | Annual | 12 | 8 | 2 | 9 | 6 | 6 |
| *Ophthalmolebias constanciae* | Annual | 16 | 6 | 3 | 5 | 3 | 20 |
| *Pachypanchax playfairii* | Non-Annual | 16 | 3 | 7 | 9 | 8 | 8 |
| *Notholebias minimus* | Annual | 12 | 2 | 4 | 5 | 2 | 4 |
| *Pituna compacta* | Annual | 11 | 1 | NA | 4 | NA | NA |
| *Pituna schindleri* | Annual | 8 | 2 | NA | 2 | 1 | NA |
| *Anablepsoides amphoreus* | Non-Annual | 11 | 3 | 2 | 9 | 3 | 15 |
| *Anablepsoides iridescens* | Non-Annual | 10 | 5 | 7 | 5 | 7 | 13 |
| *Cynodonichthys fuscolineatus* | Non-Annual | 11 | 6 | 4 | 7 | 3 | 10 |
| *Scriptaphyosemion cauveti* | Non-Annual | 13 | 5 | NA | 9 | 7 | 28 |
| *Laimosemion geayi* | Non-Annual | NA | NA | 3 | NA | NA | NA |
| *Aphyosemion gabunense* | Non-Annual | NA | NA | 2 | NA | NA | NA |
| *Pterolebias longipinnis* | Annual | NA | NA | 1 | NA | NA | NA |

**Supplementary Table 1**

Full species list and sample sizes of each species per measurement used in this study. Information on reproductive rates was sourced from Eckerström-Liedholm et al. 2017.

**Phylogeny**

In our analyses we included information on shared ancestry based on a dated phylogeny (Furness et al. 2015). We added missing taxa to the dated phylogeny by utilizing other previously published phylogenies (Murphy et al. 1999; Collier et al. 2009; Dorn et al. 2014; Ponzetto et al. 2016; Costa 2006; Costa et al. 2016; Costa et al. 2017), using the add.species.to.genus and bind.tip functions in the R package phytools (Revell 2012).

**Supplementary Table 2**

| **Parameter** | **Estimate** | **Lower CI** | **Upper CI** | **P_MCMC_** |
| --- | --- | --- | --- | --- |
| *Fixed Effects* |  |  |  |  |
| Growth | 0.03076 | 0.02667 | 0.03514 | <3*10^-04^ |
| Maturity | -1.84382 | -8.26409 | 5.37245 | 0.564 |
| Reproductive Rate | 2.91441 | 2.27411 | 3.48969 | <3*10^-04^ |
| *Random Effects* |  |  |  |  |
| Growth: Species (Growth) | 1.46*10^-04^ | 0.000079 | 2.33*10^-04^ | - |
| Maturity: Species (Growth) | 1.24*10^-01^ | 0.036453 | 2.32*10^-01^ | - |
| Reproductive Rate: Species (Growth) | 7.01*10^-03^ | -1.20*10^-03^ | 1.47*10^-02^ | - |
| Growth: Species (Maturity) | 1.24*10^-01^ | 3.65*10^-02^ | 2.32*10^-01^ | - |
| Maturity: Species (Maturity) | 3.00*10^+02^ | 1.40*10^+02^ | 5.04*10^+02^ | - |
| Reproductive Rate: Species (Maturity) | 9.53*10^+00^ | -8.11*10^-01^ | 2.14*10^+01^ | - |
| Growth: Species (Reproductive Rate) | 7.01*10^-03^ | -1.20*10^-03^ | 1.47*10^-02^ | - |
| Maturity: Species (Reproductive Rate) | 9.53*10^+00^ | -8.11*10^-01^ | 2.14*10^+01^ | - |
| Reproductive Rate: Species (Reproductive Rate) | 1.62*10^+00^ | 5.20*10^-01^ | 3.10*10^+00^ | - |
| Residual Variance (Growth) | 3.95*10^-05^ | 3.32*10^-05^ | 4.67*10^-05^ |  |
| Residual Variance (Maturity) | 1.59*10^+02^ | 1.10*10^+02^ | 2.07*10^+02^ | - |
| Residual Variance (Reproductive Rate) | 3.3*10^-01^ | 1.67*10^-01^ | 5.22*10^-01^ | - |

**Supplementary Table 2: Results of the Bayesian phylogenetic mixed model with the life-history traits, growth rate, rate of maturity and reproductive rate fitted as fixed effects. The model was run with the interactions between all possible combinations of life-history traits (growth, maturity and reproductive rate), and the life-history trait specific values for species and phylogeny, as random effects. Where, “Species” signifies the variance explained by species, “Phylogeny” signifies the variance explained by the phylogeny and “Residual Variance” is the variance that is not explained by the model. Lower and upper CIs represent 95 % credible intervals.**

**Supplementary Table 3**

| **Parameter** | **Estimate** | **Lower CI** | **Upper CI** | **P_MCMC_** |
| --- | --- | --- | --- | --- |
| *Fixed Effects* |  |  |  |  |
| (Intercept) | -0.9234833 | -1.0021684 | -0.84488 | <3*10^-04^ |
| Ontogenetic Stage (Reproducing Adult) | -0.03303 | -0.0655768 | -0.00336 | 0.0367 |
| Life-History (Non-Annual) | -0.0595241 | -0.1252026 | 0.011309 | 0.09 |
| Sex (Male) | -0.0006538 | -0.0228334 | 0.021457 | 0.9427 |
| Fish Mass (log10 transformed) | 0.8923488 | 0.8373427 | 9.50*10^-01^ | <3*10^-04^ |
| *Random Effects* |  |  |  |  |
| Species | 0.001953 | 3.94*10^-10^ | 6.63*10^-03^ |  |
| Phylogeny | 0.005978 | 2.47*10^-08^ | 1.30*10^-02^ |  |
| Residual Variance | 0.01057 | 0.00903 | 1.21*10^-02^ |  |

**Supplementary Table 3: Results of the Bayesian phylogenetic mixed model testing for sex differences in SMR, with ontogenetic stage, life-history strategy (Annual or Non-Annual) traits, sex and body mass (log10 transformed) fitted as fixed effects. The model was run with the species and phylogeny, as random effects. Where, “Species” signifies the variance explained by species, “Phylogeny” signifies the variance explained by the phylogeny and “Residual Variance” is the variance that is not explained by the model. Lower and upper CIs represent 95 % credible intervals.**

**Supplementary Table 4**

| **Parameter** | **Estimate** | **Lower CI** | **Upper CI** | **P_MCMC_** |
| --- | --- | --- | --- | --- |
| *Fixed Effects* |  |  |  |  |
| (Intercept) | 0.0997 | -0.146 | 0.292 | 0.414 |
| Ontogenetic Stage (Non-Reproducing Adult) | -0.0291 | -0.213 | 0.212 | 0.952 |
| Ontogenetic Stage (Reproducing Adult) | -0.025 | -0.247 | 0.193 | 0.839 |
| Life-History (Non-Annual) | -0.225 | -0.477 | -0.0205 | 0.0433 |
| Ontogenetic Stage (Non-Reproducing Adult): Life-History (Non-Annual) | 0.0933 | -0.138 | 0.293 | 0.461 |
| Ontogenetic Stage (Reproducing Adult:) Life-History (Non-Annual) | 0.133976 | -0.09678 | 0.366717 | 0.2407 |
| *Random Effects* |  |  |  |  |
| Ontogenetic Stage (Juvenile): Species (Juvenile) | 0.039201 | 9.67*10^-08^ | 0.08709 | - |
| Ontogenetic Stage (Non-Reproducing Adult): Species (Juvenile) | 0.013076 | -7.37*10^-03^ | 0.04272 | - |
| Ontogenetic Stage (Reproducing Adult): Species (Juvenile) | 0.004366 | -9.21*10^-03^ | 0.02556 | - |
| Ontogenetic Stage (Juvenile): Species (Non-Reproducing Adult) | 0.013076 | -7.37*10^-03^ | 0.04272 | - |
| Ontogenetic Stage (Non-Reproducing Adult): Species (Non-  Reproducing Adult) | 0.016536 | 9.01*10^-10^ | 0.0525 | - |
| Ontogenetic Stage (Reproducing Adult): Species (Non-Reproducing  Adult) | 0.00533 | -3.49*10^-03^ | 0.02387 | - |
| Ontogenetic Stage (Juvenile): Species (Reproducing Adult) | 0.004366 | -9.21*10^-03^ | 0.02556 | - |
| Ontogenetic Stage (Non-Reproducing Adult): Species (Reproducing  Adult) | 0.00533 | -3.49*10^-03^ | 0.02387 | - |
| Ontogenetic Stage (Reproducing Adult): Species (Reproducing Adult) | 0.008054 | 3.52*10^-10^ | 0.02585 | - |
| Ontogenetic Stage (Juvenile): Phylogeny (Juvenile) | 0.04304 | 1.02*10^-08^ | 0.1296 |  |
| Ontogenetic Stage (Non-Reproducing Adult): Phylogeny (Juvenile) | 0.0248 | -7.58*10^-03^ | 0.08357 | - |
| Ontogenetic Stage (Reproducing Adult): Phylogeny (Juvenile) | 0.01117 | -1.03*10^-02^ | 0.04474 | - |
| Ontogenetic Stage (Juvenile): Phylogeny (Non-Reproducing Adult) | 0.0248 | -7.58*10^-03^ | 0.08357 | - |
| Ontogenetic Stage (Non-Reproducing Adult): Phylogeny  (Non-Reproducing Adult) | 0.04519 | 1.97*10^-08^ | 0.09873 | - |
| Ontogenetic Stage (Reproducing Adult): Phylogeny (Non-Reproducing Adult) | 0.01787 | -4.63*10^-03^ | 0.0479 | - |
| Ontogenetic Stage (Juvenile): Phylogeny (Reproducing Adult) | 0.01117 | -1.03*10^-02^ | 0.04474 | - |
| Ontogenetic Stage (Non-Reproducing Adult): Phylogeny (Reproducing Adult) | 0.01787 | -4.63*10^-03^ | 0.0479 | - |
| Ontogenetic Stage (Reproducing Adult): Phylogeny (Reproducing  Adult) | 0.01821 | 4.75*10^-07^ | 0.04714 | - |
| Residual Variance | 0.05655 | 0.0498 | 0.06363 | - |

**Supplementary Table 4: Results of the Bayesian phylogenetic mixed model assessing contrasts between annual and non-annual species in metabolic rate specific to the three ontogenetic stages, in addition to species effects, with ontogenetic stages, life-history strategy and the interactions between ontogenetic stages and life-history strategy as fixed effects. The model was run with each of the possible interactions between the different ontogenetic stage specific species and phylogeny as random effects. Where, “species” signifies the variance explained by species, “phylogeny” signifies the variance explained by the phylogeny, and “residual variance” is the variance that is not explained by the mode. Lower and upper CIs represent 95 % credible intervals. The table is structured according to the standard output from R.**

**Supplementary Table 5**

| **Parameter** | **Estimate** | **Lower CI** | **Upper CI** | **P_MCMC_** |
| --- | --- | --- | --- | --- |
| *Fixed Effects* |  |  |  |  |
| Minimum Residual SMR | -0.05063 | -0.27357 | 0.15615 | 0.604 |
| Growth | 0.03014 | 0.01935 | 0.03818 | <3*10^-04^ |
| *Random Effects* |  |  |  |  |
| Minimum Residual SMR : Species (Minimum Residual SMR) | 4.17*10^-02^ | 4.72*10^-10^ | 0.100026 | - |
| Growth : Species (Minimum Residual SMR) | 3.83*10^-04^ | -9.26*10^-04^ | 0.001858 | - |
| Minimum Residual SMR : Species (Growth) | 3.83*10^-04^ | -9.26*10^-04^ | 0.001858 | - |
| Growth : Species (Growth) | 8.83*10^-05^ | 1.43*10^-10^ | 0.000194 | - |
| Minimum Residual SMR : Phylogeny (Minimum Residual SMR) | 0.057217 | 5.40*10^-09^ | 0.152747 | - |
| Growth : Phylogeny (Minimum Residual SMR) | 0.000589 | -1.07*10^-03^ | 0.002785 | - |
| Minimum Residual SMR : Phylogeny (Growth) | 0.000589 | -1.07*10^-03^ | 0.002785 | - |
| Growth : Phylogeny (Species) | 0.000106 | 5.50*10^-12^ | 0.000309 | - |
| Residual Variance (Minimum Residual SMR) | 0.062499 | 0.048716 | 0.076606 | - |
| Residual Variance (Growth) | 0.000128 | 0.000106 | 0.000152 | - |

**Supplementary Table 5: Results of the Bayesian phylogenetic mixed model assessing correlations between growth rate (in juveniles) and residual SMR, fitted as fixed effects. The model was run with the interactions between all possible combinations of growth and residual SMR specific values for species and phylogeny, as random effects. Where, “Species” signifies the variance explained by species, “Phylogeny” signifies the variance explained by the phylogeny and “Residual Variance” is the variance that is not explained by the model. Lower and upper CIs represent 95 % credible intervals.**

**Supplementary Table 6**

| **Parameter** | **Estimate** | **Lower CI** | **Upper CI** | **P_MCMC_** |
| --- | --- | --- | --- | --- |
| *Fixed Effects* |  |  |  |  |
| Minimum Residual SMR | 6.56*10^-04^ | -1.89*10^-01^ | 2.16*^-01^ | 0.981 |
| Maturity | -5.512 | -20.21 | 9.12*10^+00^ | 0.439 |
| *Random Effects* |  |  |  |  |
| Minimum Residual SMR : Species (Minimum Residual SMR) | 1.30*10^-02^ | 1.58*10^-10^ | 0.04835 |  |
| Maturity : Species (Minimum Residual SMR) | 6.94*10^-02^ | -7.44*10^-01^ | 1.04358 |  |
| Minimum Residual SMR : Species (Maturity) | 6.94*10^-02^ | -7.44*10^-01^ | 1.04358 |  |
| Maturity : Species (Maturity) | 7.04*10^+01^ | 1.21*10^-05^ | 262.866 |  |
| Minimum Residual SMR : Phylogeny (Minimum Residual SMR) | 0.06081 | 8.23*10^-06^ | 0.1155 |  |
| Maturity : Phylogeny (Minimum Residual SMR) | 1.33191 | -1.059 | 4.0172 |  |
| Minimum Residual SMR : Phylogeny (Maturity) | 1.33191 | -1.059 | 4.0172 |  |
| Maturity : Phylogeny (Species) | 318.833 | 9.57*10^-07^ | 644.309 |  |
| Residual Variance (Minimum Residual SMR) | 0.0545 | 0.04269 | 0.06742 |  |
| Residual Variance (Maturity) | 154.9912 | 99.27098 | 218.0602 |  |

**Supplementary Table 6: Results of the Bayesian phylogenetic mixed model assessing correlations between rate of maturity (in non-reproducing adults) and residual SMR, fitted as fixed effects. The model was run with the interactions between all possible combinations of growth and residual SMR specific values for species and phylogeny, as random effects. Where, “Species” signifies the variance explained by species, “Phylogeny” signifies the variance explained by the phylogeny and “Residual Variance” is the variance that is not explained by the model. Lower and upper CIs represent 95 % credible intervals.**

**Supplementary Table 7**

| **Parameter** | **Estimate** | **Lower CI** | **Upper CI** | **P_MCMC_** |
| --- | --- | --- | --- | --- |
| *Fixed Effects* |  |  |  |  |
| Minimum Residual SMR | 7.60*10^-03^ | -1.18*10^-01^ | 1.15*10^-01^ | 0.89067 |
| Reproductive rate | 2.89*10^+00^ | 1.66*10^+00^ | 3.99*10^+00^ | 0.00133 |
| *Random Effects* |  |  |  |  |
| Minimum Residual SMR : Species (Minimum Residual SMR) | 1.08*10^-02^ | 2.88*10^-11^ | 0.0297 |  |
| Reproductive rate: Species (Minimum Residual SMR) | 1.15*10^-02^ | -4.26*10^-02^ | 0.09155 |  |
| Minimum Residual SMR : Species (Reproductive rate) | 1.15*10^-02^ | -4.26*10^-02^ | 0.09155 |  |
| Reproductive rate: Species (Reproductive rate) | 5.93*10^-01^ | 9.08*10^-10^ | 2.01849 |  |
| Minimum Residual SMR : Phylogeny (Minimum Residual SMR) | 0.01413 | 2.35*10^-07^ | 0.04061 |  |
| Reproductive rate: Phylogeny (Minimum Residual SMR) | 0.03828 | -5.88*10^-02^ | 0.17212 |  |
| Minimum Residual SMR : Phylogeny (Reproductive rate) | 0.03828 | -5.88*10^-02^ | 0.17212 |  |
| Reproductive rate: Phylogeny (Species) | 1.5535 | 3.02*10^-08^ | 3.80275 |  |
| Residual Variance (Minimum Residual SMR) | 0.05602 | 0.04559 | 0.06638 |  |
| Residual Variance (Reproductive rate) | 0.33196 | 0.16149 | 0.51917 |  |

**Supplementary Table 7: Results of the Bayesian phylogenetic mixed model assessing correlations between reproductive rate (in reproducing adults, taken from Eckerström-Liedholm et al. 2017) and residual SMR, fitted as fixed effects. The model was run with the interactions between all possible combinations of growth and residual SMR specific values for species and phylogeny, as random effects. Where, “Species” signifies the variance explained by species, “Phylogeny” signifies the variance explained by the phylogeny and “Residual Variance” is the variance that is not explained by the model. Lower and upper CIs represent 95 % credible intervals.**

We thank the following people for granting us permission to use their photographs of killifish represented in Figure 3: Richard Sexton, Richard Cox, Ondřej Sedláček, Kjell Nilsson, , Anthony C. Terceira, Matheus Vieira Volcan, Marcelo Loureiro, Ivan Sazima, Frans Vermeulen ([www.itrainsfishes.net](http://www.itrainsfishes.net)), Paul V. Loiselle, Rudolf Pohlmann, Nick Neumann (on behalf of Quality Marine), Costa (2007) published in Zootaxa (Images of *Pituna compacta* and *Pituna schindleri* © Magnolia Press, reproduced with permission from copyright holder), and from Wikipedia commons (users: Cisamarc, Tommy Kronkvist, Ugau, Alexander Prokoshev, Andrew Bogott and Gustavo Grandjean.
